# Harmonic decomposition of spacetime (HADES) framework characterises the spacetime hierarchy of the DMT brain state

**DOI:** 10.1101/2023.08.20.554019

**Authors:** Jakub Vohryzek, Joana Cabral, Christopher Timmermann, Selen Atasoy, Leor Roseman, David J Nutt, Robin L Carhart-Harris, Gustavo Deco, Morten L Kringelbach

**Affiliations:** Centre for Eudaimonia and Human Flourishing, Linacre College, University of Oxford; Department of Psychiatry, University of Oxford, Oxford, United Kingdom; Center for Music in the Brain, Aarhus University, Aarhus, Denmark; Center for Brain and Cognition, Computational Neuroscience Group, Department of Information and Communication Technologies, Universitat Pompeu Fabra, Barcelona, Spain; Life and Health Sciences Research Institute, School of Medicine, University of Minho; ICVS/3B’s - PT Government Associate Laboratory, Braga/Guimarães, Portugal; Centre for Psychedelic Research, Department of Brain Sciences, Imperial College London, London, United Kingdom; Departments of Neurology and Psychiatry, University of California San Francisco, US; Institució Catalana de la Recerca i Estudis Avançats (ICREA), Barcelona, Spain; Department of Neuropsychology, Max Planck Institute for Human Cognitive and Brain Sciences, Leipzig, Germany; School of Psychological Sciences, Monash University, Melbourne, Australia

**Keywords:** Spatio-temporal Brain Dynamics, DMT, Harmonic Modes

## Abstract

The human brain is a complex system, whose activity exhibits flexible and continuous reorganisation across space and time. The decomposition of whole-brain recordings into harmonic modes has revealed a repertoire of gradient-like activity patterns associated with distinct brain functions. However, the way these activity patterns are expressed over time with their changes in various brain states remains unclear. In this study, we develop the Harmonic Decomposition of Spacetime (HADES) framework that characterises how different harmonic modes defined in *space* are expressed over *time*, and, as a proof-of-principle, demonstrate the sensitivity and robustness of this approach to specific changes induced by the serotonergic psychedelic N,N-Dimethyltryptamine (DMT) in healthy participants. HADES demonstrates significant decreases in contributions across most low-frequency harmonic modes in the DMT-induced brain state. When normalizing the contributions by condition (DMT and non-DMT), we detect a decrease specifically in the second functional harmonic, which represents the uni- to transmodal functional hierarchy of the brain, supporting the hypothesis that functional hierarchy is changed in psychedelics. Moreover, HADES’ dynamic spacetime measures of fractional occupancy, life time and latent space provide a precise description of the significant changes of the spacetime hierarchical organization of brain activity in the psychedelic state.

## Introduction

The brain is endowed with complex dynamics and can be perceived along spatial and temporal dimensions [1]. Traditionally, neuroscience has focused on delineating and studying localised cortical regions to map brain function in a temporarily static fashion [2]. However, recent developments in neuroscience have started to indicate more spatially continuous representations of functional topography [3], [4], and at the same time to stress the importance of temporally varying brain dynamics [5]. Despite such progress, it remains unknown what underlying mechanisms drive, on one hand, the gradient-like organisation of cortical topography, and on the other, the waning and waxing of the brain’s spatiotemporal patterns of activity.

Here, we propose Harmonic Decomposition of Spacetime (HADES) as a new model of hierarchical processing across both spatial and temporal dimensions. Historically, Brodmann’s interactive atlas of cellular morphology and organisation has given rise to the view of functional specialisation of individual brain areas [6], [7]. Spatially, this suggests a sharp delineation between cortical areas in terms of their anatomy and function. However, supported by evolutionary and developmental neuroscience [8], [9], cortical gradients have challenged this view by suggesting gradually varying boundaries between and within brain regions, both in terms of function and anatomy [3], [4], [10]. Functionally, gradient-like organisation proposes an intrinsic coordinate system of human brain organisation continuously varying from unimodal to transmodal cortical areas [3], [11]. Similarly, topographical maps of retinotopy, somatotopy and tonotopy have shown smooth variation of anatomy and function within brain areas [12]–[15].

Along the temporal dimension, studies of dynamic functional connectivity in fMRI have revealed the importance of characterising the temporal features of brain activity as opposed to the static picture described by known resting-state networks [5], [16]. Such approaches describe temporal functional connectivity in terms of sliding-window analysis [17], by considering the most salient events in the timeseries [18], [19] constrained by structural connectivity [20], [21], as a temporal process of hidden states [22], [23] or as a temporal trajectory in a landscape of attractors [24], [25]. Broadly, these approaches share the description of complex brain dynamics in terms of spatial patterns expressed in time and therefore can be represented in terms of the patterns’ fractional occupancy, life times or probability of transitions.

Here, HADES characterizes brain’s spatio-temporal activity in terms of harmonic modes defined in *space* and expressed over *time*. For that end, we derived the functional harmonics (FHs) [4] and their temporal expression by decomposing fMRI data into functional harmonics via harmonic decomposition [26]. The motivation for HADES is, on one hand, to account for an increasing spatial scale from neuronal circuits to large-scale brain networks, and on the other, for its temporal evolution. Furthermore, HADES attempts to improve on the earlier methods limitations demonstrating spatial interpretability, modelling feasibility and analysis flexibility [27], [28]

One of the most potent psychedelic (i.e. ‘mind-manifesting’) experiences is induced by the N,N - Dimethyltryptamine (DMT) - a naturally occurring serotonergic psychedelic [29]. Unlike psilocybin and LSD, its expression is marked by a short duration of the psychedelic experience. It is often associated with alterations in visual and somatic effects. At high doses, a complete dissociation from the external environment precedes an immersion into mental worlds or dimensions described as "other" but not less "real" than the one inhabited in normal waking consciousness. Such experiences correlate with subjective rating items such as "I experienced a different reality or dimension", "I saw geometric patterns" and "I felt unusual bodily sensations" [30], [31]. It is these qualities of one’s conscious experience that motivate a renewed interest in DMT drawing parallels with phenomena such as the near-death experience (NDE) and dreaming [32].

Furthermore, like other psychedelics, DMT may have clinical relevance and is currently being trialled for the treatment of depressive symptoms [33], [34]. Studies with Ayahuasca, containing DMT itself as well as monoamine oxidase inhibitors (MAOIs), have shown promising results in patients with depression [35]. However, further investigations exploring the neural and plasticity dynamics of DMT experiences are necessary to provide mechanistic accounts for the relevance of DMT and related psychedelics for the treatment of mental health disorders [36]–[38].

In the brain, psychedelics enhance the richness of spatio-temporal dynamics along both the temporal and spatial dimensions. This has been corroborated by repertoire broadening of functional states and increases in temporal complexity as well as shifting of the brain to a more integrated state with the subversion of functional systems [39]–[42]. Consistently, neuroimaging with DMT has revealed an increase in global functional connectivity – featuring a functional network disintegration and desegregation that is reliable feature of the psychedelic state, and a collapse of the unimodal to transmodal functional gradient [31]. Taken all together, the current findings and subjective reports are in line with the entropic [43], [44] and anarchic brain [45] models, where an increase in entropy of spontaneous brain activity parallels the undermining of hierarchically organised brain function [43]–[45].

Here we use fMRI data from the DMT-induced state to describe HADES’s multifaceted applications. Empirically, based on anarchic brain or ‘Relaxed Beliefs Under Psychedelics’ (REBUS) model, as well as findings of enhanced signatures of criticality under these compounds [26], [41], [46], we hypothesised that the DMT state is associated with a flatter hierarchy of cortical functional organisation with enhanced integrative properties across the cortex.

## Results

Harmonic Decomposition of Spacetime (HADES) describes the spatio-temporal dynamics in terms of spatial bases (defined from the brain’s communication structure) and the spatial bases functional contributions to the fMRI recording evolving in time. To do so, we first constructed dense functional connectome from the Human Connectome Project (HCP) S1200 release of 812 subjects (**Figure 1B**). The dense functional connectome was represented as a sparse, symmetric, and binary adjacency matrix (**Figure 1C**) and decomposed into the functional harmonics (ψ_k_(x)) using the eigen-decomposition of the graph Laplacian applied to the dense functional connectome (**Figure 1D**). Consistent with [4], we focused our analysis on the first 11 lowest functional harmonics together with the global zeroth harmonic. We analysed functional significance of the functional harmonics by comparing them to the Yeo seven and seventeen functional networks (**Figure SI1**). To obtain the temporal signature, we further projected the individual harmonics on the fMRI timeseries (in surface representation), using functional harmonic decomposition, and thus calculated the FHs temporal weights (**Figure 1E**). We reconstructed the timeseries with a few harmonics to motivate the similarity to the empirical data (**Figure SI2**). Then, using a collection of non-dynamic and dynamic measures (**Figure 1F and 1G**) and latent space representation (**Figure 1H**), we applied HADES to show its viability in researching rich and complex brain dynamics in different brain states and illustrate this in the context of the DMT-induced state.

**Figure 1.**
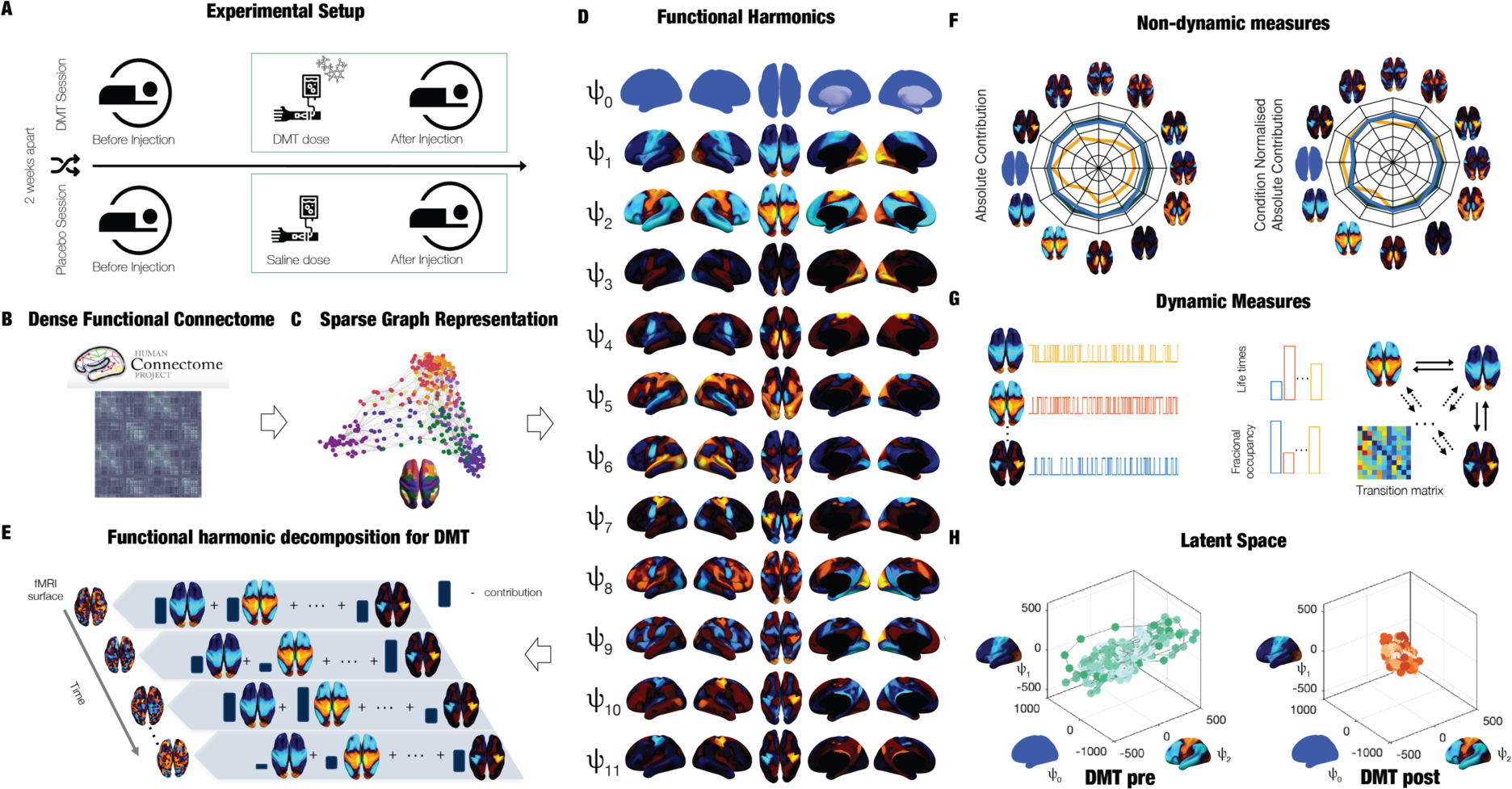
Overview of HArmonic DEcomposition of Spacetime (HADES) framework. **A)** Here we used HADES to analyse data from DMT-induced resting-state fMRI in healthy participants and show the design for this experiment. **B)** HADES uses the dense functional connectome constructed from the HCP S1200 release of 812 subjects to **C)** construct a graph representation as a sparse, symmetric, and binary adjacency matrix of the dense functional connectome. **D)** First, Functional Harmonics (*ψ_k_*(*x*)) are obtained from the Laplacian decomposition of the sparse adjacency matrix. **E)** Functional harmonic decomposition is computed by projecting individual harmonics on the fMRI timeseries (surface representation) and calculating their contributions. **F)** From this decomposition, HADES can be used to compute non-dynamic measures for the first 12 Functional Harmonics – Absolute Contribution and Condition Normalised Absolute Contribution on any neuroimaging dataset. **G)** Importantly, HADES can also be used to construct dynamic measures for the first 12 Functional Harmonics – Fractional Occupancy, Life Times and Transition Matrix. **H)** These can be measures can be used as latent space representation as the temporal trajectory embedded in the Functional Harmonics space.

### Absolute Contribution across Functional Harmonics

To quantify contributions of individual harmonics in the different conditions, we computed the absolute and condition-normalised absolute contributions of each harmonic (**Figure 2A**). The absolute contribution results show a decrease in the DMT-induced state (compared to DMT before injection and placebo-induced states) across most of the 11 FHs except of the global FH (green star: p-value < 0.05 Bonferroni-corrected paired t-test, red star: p-value < 0.05 uncorrected paired t-test). This is contrasted by the condition-normalised absolute contribution results demonstrating an increase in the global FH and a decrease in FH 2 after DMT injection versus before injection and the placebo data (green star: p value < 0.05 Bonferroni-corrected paired t-test, red star: p-value < 0.05 uncorrected paired t-test, **Figure 2B**). Spider plots in **Figure 2A** and **2B** represent a visual redistribution of FHs across different conditions for the two measures.

**Figure 2:**
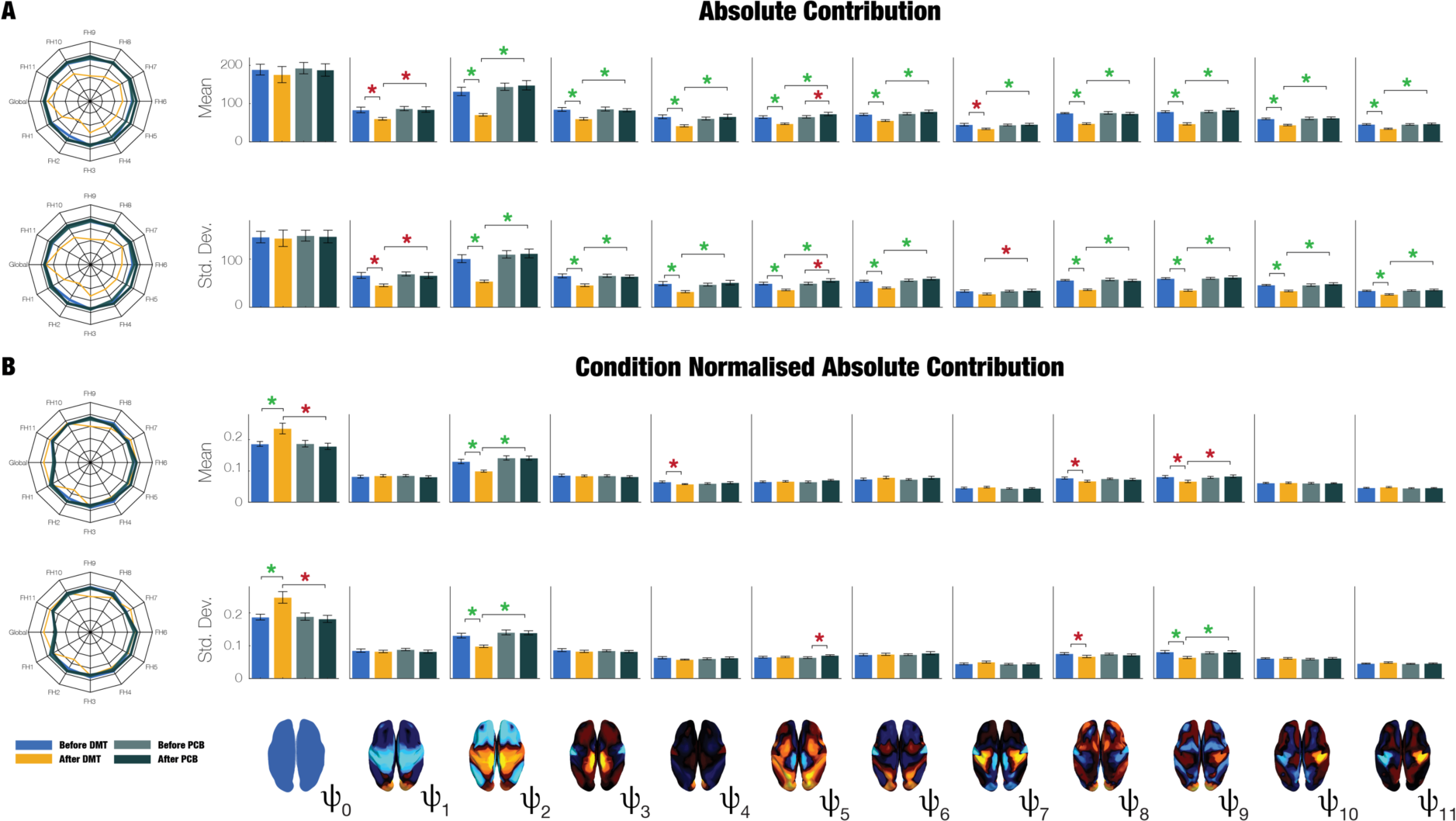
Harmonic Spatial Analysis of DMT and placebo neuroimaging data. The harmonic spatial analysis of the neuroimaging data shows that the contribution of Functional Harmonic *ψ*_"_ (FH*ψ*_"_) is very significantly reduced (p<0.05, Bonferroni corrected) when participants were given DMT, both in terms of absolute and normalised contribution. **A)** Specifically, the absolute contribution across the first 12 FHs is shown both visually, on a spider plot, and statistically for individual FH across the four DMT-based conditions. The results show a decrease in the DMT-induced state (compared to DMT before injection and the placebo state) across many of the 12 FHs except the global FH *ψ*_#_ (green star p-value < 0.05 Bonferroni corrected paired t-test, red star p-value < 0.05 not Bonferroni corrected paired t-test). **B)** Equally, we show the Normalised Absolute Contribution across the first 12 FHs represented both visually, on a spider plot, and statistically for individual FHs across the four DMT-based conditions. Again, the results demonstrate an increase in the global FH *ψ*_#_but specifically a decrease in FH *ψ*_"_ compared to DMT before injection and the placebo state (green star p-value < 0.05 Bonferroni corrected paired t-test, red star p-value < 0.05 not Bonferroni corrected paired t-test).

### Dynamic Measures of HADES

To assess the temporal evolution of FH weights, we apply a winner-takes-all approach whereby we select the most prominent FH at every time point and compute Fractional Occupancy (FO) and Life Times (LT) of each FH. In **Figure 3A** and **B**, we show results when choosing the 11 FHs. We excluded the zeroth FH in this analysis to focus on the dynamical properties of functionally resolved FHs. As before, strongest statistical significance for FO and LT is observed in *ψ*_2_ (green star: p value < 0.05/(# of FH) paired t-test, red star: p-value < 0.05 uncorrected paired t-test, **Figure 3C**). Furthermore, we computed the first order Markov process in terms of the Transition Probability Matrix (TPM) (**Figure SI 3A**). We report statistics for the two DMT conditions (p-value < 0.05 uncorrected paired t-test).

**Figure 3.**
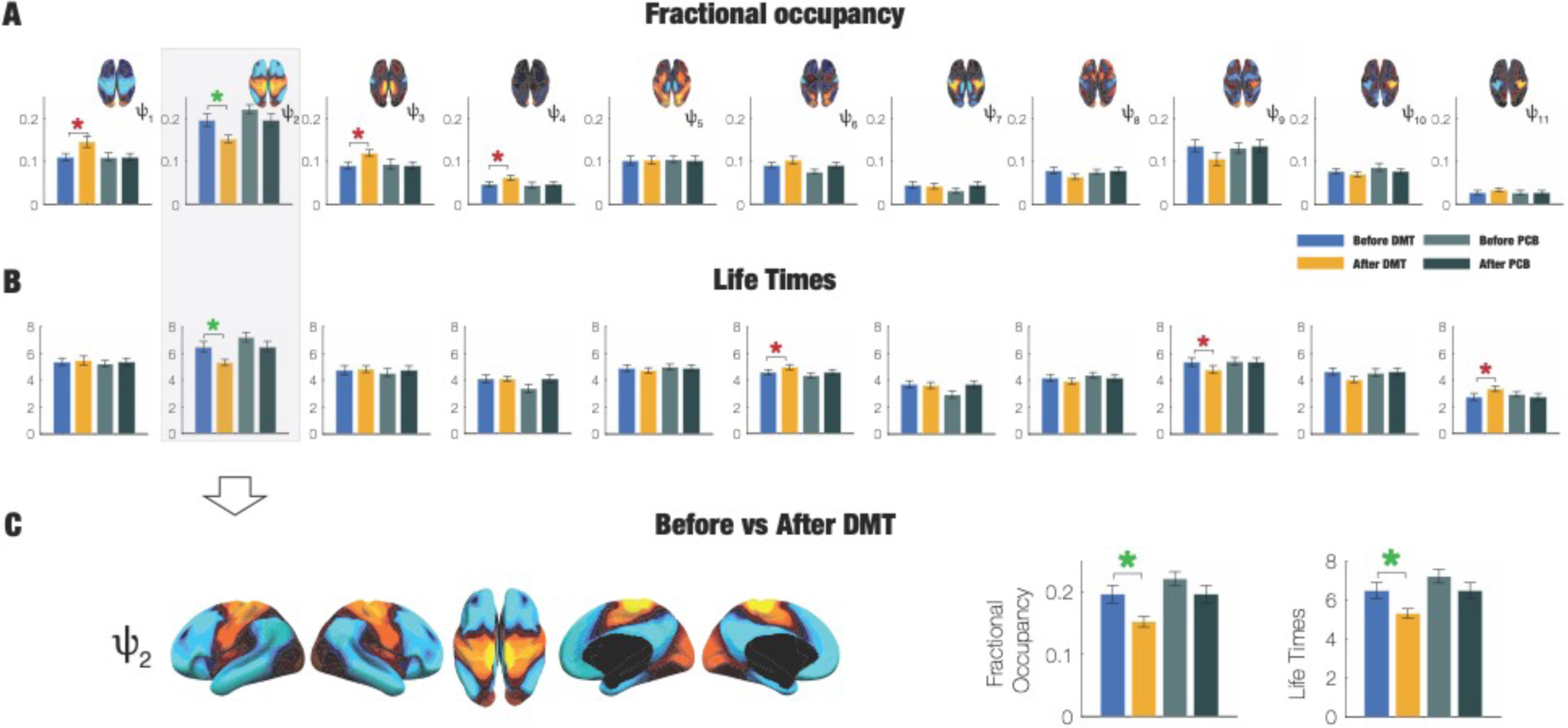
Spatiotemporal HADES analysis for the 11 Functional Harmonics (FH) Extending the spatial analysis into the spatiotemporal domain again shows that Functional Harmonic *ψ*_"_ (FH*ψ*_"_) is significantly reduced in the DMT condition. **A)** Specifically, Fractional Occupancy was found to be statistically different in the *ψ*_"_. **B)** Life Times were found statistically different in the *ψ*_"_ (green star: p value < 0.05 (# of *ψ*_$_) where n=11 paired t-test, red star: p-value < 0.05 uncorrected paired t-test). **C)** The full spatial extent of FH *ψ*_"_ is shown along with the significant results for Fractional Occupancy and Life Times.

### Latent Space

Functional harmonics were used as the basis of a latent space representation in which the temporal trajectory of the brain dynamics was embedde in the latent space representation of the 12 FHs (**Figure 4A**, here visualised for the first three FHs with colour shading representing the temporal trajectory). To further analyse how the temporal embedding in this latent space changes, we defined the expansion/contraction of the trajectory in term of the latent dimension spread. The DMT-induced state contracts the contribution of the FHs across the board. Latent dimension spread was computed for all the 12 FHs i.e., 12^th^ dimensional space for the four conditions. We also report its statistics (green star p-value < 0.05 Bonferroni corrected paired t-test). The temporal trajectory significantly contracts in the DMT-induced state.

**Figure 4.**
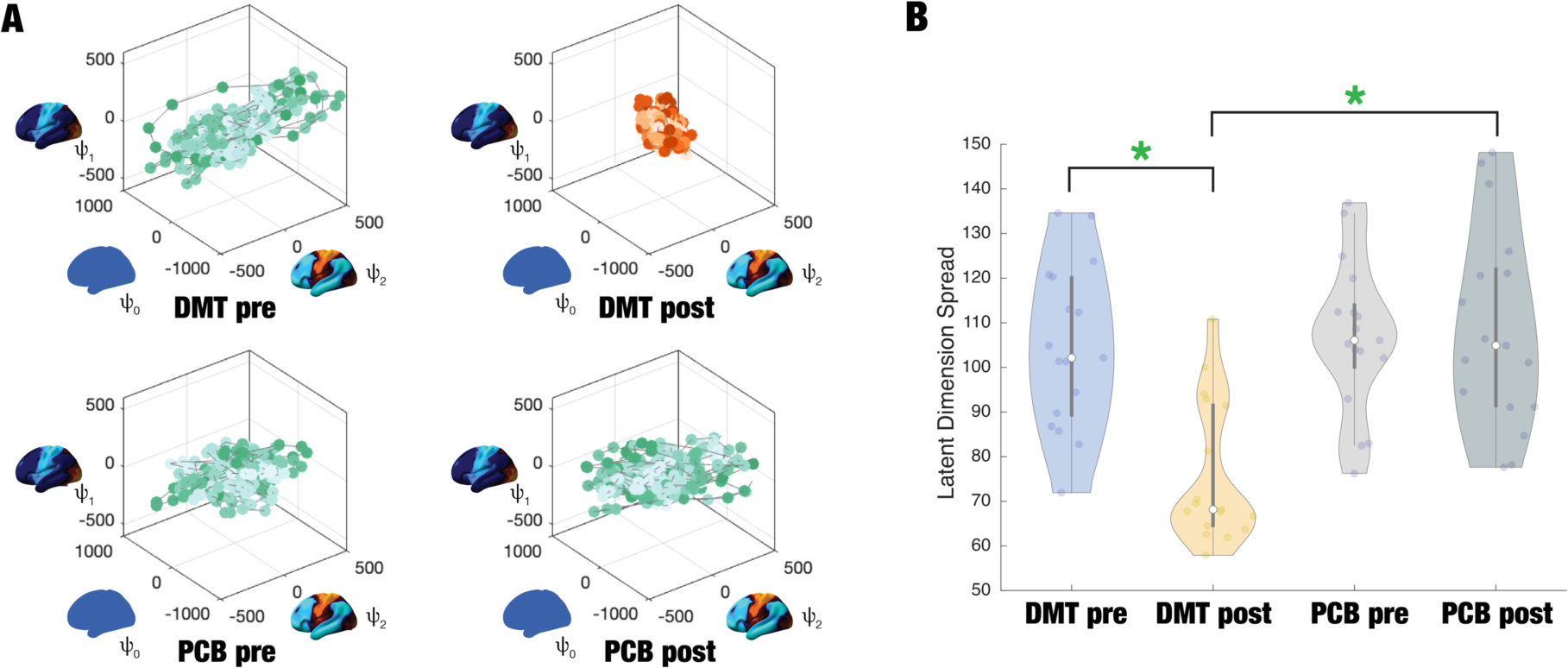
Latent Space Representation of neuroimaging data using the 12 Functional Harmonics (FHs) Importantly, HADES can be used to create a latent space representation of the DMT neuroimaging data that immediately brings out important spacetime differences. **A)** Here we show the figures with Latent Space Representation using the first three FHs for visualisation of the neuroimaging data. The green colour shading represents the temporal trajectory embedded in the three latent spatial dimensions of the FHs of DMT_pre, PCB_pre and PCB_post. As can be immediately seen for the DMT-induced state (DMT_post) there is a clear contraction of the contribution of the FHs across board (shown in red colour shading). **B)** This can be directly quantified in terms of the Latent Dimension Spread computed for all the 12 FHs i.e. 12^th^ dimensional space for the four conditions. As can be see DMT_post is significantly different from DMT_pre and PCB_post (green star p-value < 0.05 Bonferroni corrected paired t-test).

## Discussion

In this study, we describe our novel HArmonic DEcomposition of Spacetime (HADES) framework. HADES is designed to be a sensitive and precise measure of the spacetime features of neuroimaging data. The framework uses the first 12 functional harmonics associated with the lowest spatial frequencies derived from the dense functional connectome of the brain from a large group of 812 healthy participants. Any neuroimaging data can then be decomposed in terms of the spacetime contributions of these functional harmonics. Here, as proof-of-principle, we used HADES to analyse the DMT-induced brain state in healthy participants and found a significant change of brain hierarchy in line with theoretical predictions of the anarchic brain hypothesis, also known as ‘REBUS’ [45].

Consistent with previous literature, we have demonstrated the functional relevance of functional harmonics [4]. Moreover, we have demonstrated that an empirical fMRI signal can be accurately reconstructed with a subset of functional harmonics. Applying HADES to the DMT-induced state has shown decreases in absolute contribution across most FHs, while the global FH has remained unchanged. However, when looking at condition-normalised absolute contribution in individual subjects, a decrease in FH *ψ*_2_ was mirrored by an increase in the global harmonic. These results motivate a non-trivial reconfiguration whereby the DMT-induced state decreases in overall magnitude with a relative increase towards the global substate and a decrease of FH *ψ*_2_ representative of the functional hierarchies of the brain. This was further reinforced by the analysis of functional harmonic dynamics with decreases both in fractional occupancy and lifetimes of FH *ψ*_2_ demonstrating further dynamic collapse of this harmonic. Lastly, when the temporal trajectories were embedded in the latent space of the functional harmonic, the DMT-induced state showed significant contraction of its temporal trajectory spread.

Remarkably, FH *ψ*_2_ rresembles the so-called ‘principal gradient’ - i.e., a unimodal to transmodal gradient previously found to explain the greatest proportion of variance in a principal components analysis of cortical functional connectivity [3]. This gradient has been proposed to reflect a hierarchy of brain function from low- to high-order cognitive networks We have argued that psychedelic-induced states result in the undermining of functional systems’ hierarchies in the brain as proposed and experimentally corroborated by the model known as ‘REBUS and the anarchic brain’ [31], [45], [47]. Furthermore, the relative increase in global FH speaks to a less functionally defined and more integrated global substate under the influence of DMT. Indeed, on the RSN level, psychedelic-induced states have been shown to subvert within functional network-connectivity, especially in higher-order fronto-parietal and default mode networks [31], [42], [48], [49], while enhancing between-network connectivity and overall global and integrative tendencies [31], [39].

Traditionally, neuroscience has focused on delineating and studying localised cortical regions to map the brain’s function. Such approach has been of importance albeit with fragmented insights as to how multiscale brain organisation gives rise to complex spatio-temporal dynamics and ultimately behaviour. A recent development in system neuroscience has been that of cortical gradients [3]. This proposes an intrinsic coordinate system of human brain organisation continuously varying from unimodal to transmodal cortical areas [11]. Gradient-type organisation has been demonstrated in terms of myelination [50], anatomical structure [10], white matter tract length [51], evolutionary expansion [52], ontogenetic expansion [53], temporal processing [54], semantic processing [55] and physiologically coupled travelling waves [56]. The framework of multidimensional harmonic representation and decomposition [4], [26], [57] adds to this list by decomposing brain activity maps into frequency-specific communication channels that unveil contributions of connectivity gradients and cortical parcellations to brain function. HADES extends these frameworks by considering the dynamic aspects of these frequency-specific channels of functional communication.

The brain as a complex system is hypothesised to manifest hierarchies across time and space. Indeed, such a nested organisation was suggested both in terms of the structural architecture of the brain as well as its temporal frequencies [58], [59]. Functional harmonics are by construction intrinsically ordered according to their spatial frequencies and as such provide a multiscale representation of brain activity across cortical space. Intuitively, spatial frequencies relate to temporal frequencies of oscillations and therefore further research with modalities such as EEG or MEG will be interesting for drawing a closer relationship between the two [40].

Previously, connectome harmonics have been used to decompose the brain’s spatio-temporal activity into a combination of time-varying contributions [26]. Using long-range and local connectivity as an underlying structure has been relevant in exploring the structure-function relationship of large-scale brain organisation [57]. However, it seems that structural connectivity alone cannot explain the emergence of rich and spontaneous activity of the human brain [60], [61]. Firstly, neocortex is endowed with remarkable heterogeneity in cytoarchitecture. This will result in various computational differentiation across the cortex, for example in terms of temporal processing [54]. Secondly, the neuromodulatory system is known to alter the electrical composition of neurons and thus exercise non-linear effects on the emergent activity of various microcircuits across the brain [62], [63]. The hypothesis here is that the communication structure of dense FC has implicitly embedded within it information on anatomical structure, cortical computational heterogeneity as well as neuromodulatory expression and as such serves as a prominent candidate to be used for the derivation of fundamental functional building blocks of spatiotemporal activity [4]. This in turn is expanded upon in the HADES framework with dynamic measures and latent space embeddings, whereby the emphasis is on the importance of the temporal dimension along which these spatio-temporal blocks building unfold.

Latent space representation has become an important research topic in neuroscience due to its ability to retrieve meaningful features contained in large and complex datasets [64]. It is possible to identify patterns and relationships in a lower-dimensional space between regions and between cognitive processes as the underlying computations giving rise to cognitive functions are likely to be integrated [1]. There are many techniques that serve this purpose from more traditional linear approaches such as singular value decomposition or principal component analysis [65], to popular techniques based on independent component analysis [66]. More recent works use autoencoders as an elegant way in compressing fMRI signal while accounting for non-linearity in the data [67]. Here, we chose functional harmonics as they preserve nonlinear relationship between regions, and have multiscale and interpretable representation of its latent dimensions [4], [68]. However, it is to be noted that the idea of HADES as a framework span beyond the actual representation of the dimension of the latent space (here in terms of functional harmonics) as it attempts to combine the spatial and temporal representation of the complex brain dynamics. Moreover, in theory, other techniques could be applied in a similar way as to account for the complex spatio-temporal activity of the human brain.

A limitation of the current approach for describing functional harmonics propagating in time is that it might be too reductionist. ’Winner-takes-all’ is a powerful technique summarising the brain’s dynamics in terms of fractional occupancy and lifetimes of the functional harmonics. However, it considers only one FH to be active at a given timepoint and as such might neglect other potential important information included in other FHs. Future work should implement weighted contributions of individual FHs at given timepoints and as such more completely describe the multidimensional representation of spatio-temporal dynamics.

## Conclusion

Taken all together, in this study we have introduced a new method called Harmonic Decomposition of Spacetime (HADES) to describe spatio-temporal dynamics of the brain. Using Functional Harmonics (FHs) derived from the brain’s communication structure, HADES models dynamics as weighted contributions of FHs evolving in time. Firstly, we verified the functional relevance of FHs with known resting-state networks showing both gradient-like and network-based organisation. Then, we reconstructed aspects of the original timeseries with only 100 FHs and their contributions. Furthermore, we applied HADES to the DMT-induced state. We showed how condition-normalised and absolute contributions can be used to demonstrate suppression of functional hierarchy and enhancement of whole brain integration. Lastly, we demonstrated similar findings of impaired hierarchical organisation in dynamic terms as shown by fractional occupancy and life times of FH *ψ*_2_. These findings corroborate the REBUS and anarchic brain model of psychedelic action by demonstrating dynamic changes to brain functional hierarchies.

## Supporting information

HADES - Supplementary Information

## Acknowledgements and Fundings

Jakub Vohryzek is supported by EU H2020 FET Proactive project Neurotwin grant agreement no. 101017716, Morten L. Kringelbach is supported by the European Research Council Consolidator Grant: CAREGIVING (615539), Pettit Foundation, Carlsberg Foundation and Center for Music in the Brain, funded by the Danish National Research Foundation (DNRF117). Joana Cabral is supported by ”la Caixa” Foundation LCF/BQ/PR22/11920014, Spain and by the Portuguese Foundation for Science and Technology UIDB/50026/2020 and UIDP/50026/2020, Portugal. Gustavo Deco is supported by the Spanish Research Project PSI2016-75688-P (Agencia Estatal de Investigación/Fondo Europeo de Desarrollo Regional, European Union); by the European Union’s Horizon 2020 Research and Innovation Programme under Grant Agreements 720270 (Human Brain Project [HBP] SGA1) and 785907 (HBP SGA2); and by the Catalan Agency for Management of University and Research Grants Programme 2017 SGR 1545.

## Material and methods

### Experimental Data

#### HCP Functional MRI

The dataset used for the analysis was made publicly available by the Human Connectome Project (HCP), WU-Minn Consortium (Principal Investigators: David Van Essen and Kamil Ugurbil: 1U54MH091657). This project was made possible by funding from the sixteen NIH Institutes and Centres supporting the NIH Blueprint for Neuroscience Research; and by the McDonell Centre for Systems Neuroscience at Washington University.

#### Dense Functional Connectome

To define the appropriate functional basis, we used the dense functional connectome as part of the HCP 1200 Subject Release. The data is freely downloadable (with a connectomeDB account) at https://db.humanconnectome.org under the zip-file called 812 Subjects, recon r227,

Dense Connectome. Details about the dense functional connectome pipeline can be found on the same website under the following pdf ‘HCP1200-DenseConnectome +PTN+Appendix-July2017.pdf’. In brief, out of the 1200 HCP subjects, 1003 have undergone four rsfMRI runs (total of 4800 timepoints). An improved reconstruction software (’recon2’) was used on a further subset of 812 participants. Timeseries were minimally processed, had artefacts removed with ICA+FIX and were inter-subject registered. Further group-PCA was performed on the temporally demeaned and variance normalised timeseries. The outputs of the group-PCA are used to create the dense connectome. This can be thought of as a low-noise regularised equivalent of concatenating individual subject’s gray-ordinate timeseries and calculating the correlation between all the individual grey-ordinate timeseries, to create a dense functional connectome (**Figure 1A**).

#### DMT dataset

The complete description of the participants, experimental design and acquisitions parameters can be found in [30], [31]. A group of 25 participants was recruited in a single-blind, placebo-controlled, and counter-balanced design. Subjects were considered for the study unless they were younger than 18 years of age, lacked experience with a psychedelic, had a previous negative response to a psychedelic and/or currently suffered from or had a history of psychiatric or physical illness. Out of the 25 participants, 20 completed the whole study (7 female, mean age = 33.5 years, SD = 7.9). A further 3 subjects were excluded due to excessive motion during the 8 minutes DMT recording (more than 15% of volumes scrubbed with framewise displacement (FD) of 0.4 mm).

#### Experimental Paradigm

In total, all subjects were scanned on two days, two weeks apart, each consisting of two scanning sessions. The initial scan lasted 28 minutes with the 8th minute marking the intravenous administration of either DMT or placebo (saline) (50/50 DMT/placebo). Subjects were asked to lay in the scanner with their eyes closed (wearing an eye-mask). After the recording, assessment of subjective effects was carried out. The second session was identical to the first except for the assessment of subjective intensity scores at every minute of the recording. The experimental design also included simultaneous EEG recording during the sessions (**Figure 1A**).

#### Acquisition Parameters

The experiment was performed on a 3T scanner (Siemens Magnetom Verio syngo MR 12) with compatibility for EEG recording. A T2 -weighted echo planar sequence was used. In brief, the parameters were as follows: TR/TE = 2000ms/30ms, acquisition time = 28.06 minutes, flip angle = 80o, voxel size = 3×3×3 mm^3^ and 35 slices with 0 mm interslice distance. T1-weighted structural scans of the brain were also acquired.

#### fMRI Pre-processing

For fMRI pre-processing, a pipeline previously developed for an LSD experiment was used, which can be accessed in the supplementary information of [48]. Briefly, the following steps were applied 1) despiking, 2) slice-timing correction, 3) motion correction, 4) brain extraction, 5) rigid body registration to structural scans, 6) non-linear registration to 2mm MNI brain, 7) motion-correction scrubbing, 8) spatial-smoothing (FWHM) of 6 mm, 9) bandpass filtering into the frequency range 0.01-0.08 Hz, 10) linear and quadratic detrending, 11) regression of 9 nuisance regressors (3 translations, 3 rotations and 3 anatomical signals). Lastly, the timeseries were projected from MNI voxel-space to the HCP surface vertex-space using the HCP command -volume-to-surface-mapping.

#### Functional Harmonics

Functional harmonics are described by the eigenvectors of the Laplacian applied to a graph representation of the human brain’s communication structure [4]. This graph is constructed as a binarization of the dense functional connectome ℜ = (*v*, *ε*), where each node, *v* = {*x*_*i*_| ∈ 1, …, *n*}, corresponds to one of the n = 59 412 brain vertices and, for each node/vertex n, an edge, *ε* = {*e*_*ij*_| ∈ *v* × *v*}, is defined to the 300 most correlated vertices, according to the correlation values from the original dense functional connectome (**Figure 1B**). Then, the resulting graph is thus a sparse, symmetric, and binary adjacency matrix (**Figure 1C**) as follows,

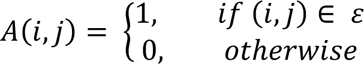

Then, the discrete counterpart of the Laplace operator, Δ, is applied to the adjacency matrix A in the following manner,

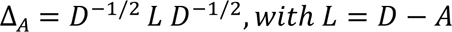

where D is the diagonal degree matrix, 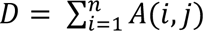. Lastly, Functional Harmonics, *ψ^k^*(*x*_*i*_), *k* ∈ 1, …, *n* were computed as eigenvectors of the following eigenvalue problem,

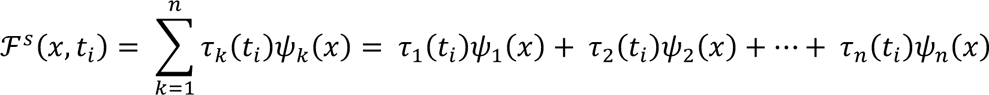

where *λ_k_*, *k* ∈ 1, …, *n* are the associated eigenvalues of Δ_*A*_ (**Figure 1D**).

#### Functional Harmonic Decomposition

To describe how Functional Harmonics evolve in time, we weighted their contribution, *τ*, for each participant at every timepoint, *t*, of the recording *F^s^*(*x*, *t*), and thus, retrieved timecourses of individual harmonic contributions (**Figure 1D**) in the following format,

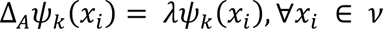

where *τ_k_* is the contribution of the *k^th^* Functional Harmonic *ψ_k_*(*x*) to the fMRI recording *F^s^*(*x*, *t_i_*) at time *t_i_*. Formally, the Functional Harmonic contributions are described as *τ_k_*(*t*) = 〈*F^s^*(*x*, *t*), *ψ_k_*〉 (**Figure 1E**).

#### Non-dynamic Measures

Functional Harmonic contribution *τ_k_*(*t*) at each timepoint *t* represents the weight of a given Functional Harmonic *ψ_k_*(*x*) at that particular fMRI timepoint, *F^R^*(*x*, *t_i_*). Its absolute value can be defined as the absolute contribution as follows: *P*(*ψ*(*x*), *t*) = |*τ_k_*(*t*)|. Here, we further define the mean absolute and condition-normalised absolute contribution as the time-averaged overall absolute contribution of each harmonic, and as the time-averaged condition-normalised absolute contribution by the sum of all the Functional Harmonic magnitudes of each participant and condition, respectively. In other words, absolute contribution describes the overall state of each Functional Harmonic for every participant and condition, and condition-normalised absolute contribution depicts the relative redistribution for a given Functional Harmonic in relationship to the rest of the Functional Harmonics (**Figure 1F**).

#### Dynamic Measures

To summarise dynamics of Functional Harmonics, we chose to describe each timepoint by its dominant Functional Harmonics, i.e., a Functional Harmonic with the largest contribution at a given timepoint. As such, we were able to depict the individual timeseries as a sequence of dominant Functional Harmonic contributions. we further defined Fractional Occupancy, Life Times and Transition matrix as the probability of a given Functional Harmonic being active during the duration of the recording, the averaged consecutive period a given Functional Harmonic was on, and first order Markov-chain for the Functional Harmonics respectively (**Figure 1G**).

#### Latent Space

Latent space serves as a lower-dimensional representation of high-dimensional data. Here, we have used the spatial patterns, described by Functional Harmonics, to embed the temporal activity in N-dimensional space where N is the number of FHs. As such it is possible to quantify the changes in temporal dynamics of FHs. Here, we define measure of Latent Dimension Spread that quantifies the amount of temporal trajectory expansion or contraction. It is defined as the average of the 11 FHs of the standard deviation of the Functional Harmonic contribution *τ_k_*(*t*) over time (**Figure 1H**).

