## Supplementary material for "Harmonic decomposition of spacetime (HADES) framework characterises the spacetime hierarchy of the DMT brain state": HADES - Supplementary Information

### Parcellation and Reference Functional Networks

Functional Harmonics in vertex dimensions ($n = 59,412$) were reduced to the Schaefer multiscale atlas of varying numbers of brain regions. Schaefer400 was applied in the main text to compare FH with the known canonical networks (7 and 17 Yeo resting-state networks [1]) but other scales were also considered (Schaefer100, 200, 300, 500 and 1000). The parcellated data was obtained by averaging vertices belonging to a given brain region as defined by the Schaefer parcellation. Unlike standard volumetric fMRI to atlas registration, the parcellated data were obtained in the surface space of 59, 412 vertices. I used the following Schaefer HCP surface templates www.github.com/ThomasYeoLab.

**Functional Relevance**

Firstly, we compute and verify the overlap of Functional Harmonics with the canonical networks - specifically the 7 and 17 resting-state networks as described in the following study [1]. Previously, Glomb et al. have demonstrated the neurophysiological relevance of Functional Harmonics [2]. In **Figure SI 1A**, we display Functional Harmonics as computed both by HADES and by [2] for visual inspection. Apart from the Functional Harmonic 10, the patterns are consistent across the two datasets. As the calculation steps were kept identical, a possible discrepancy might have arisen from the dense functional connectome itself where an improved reconstruction software was used for the HADES dataset (in HADES: S1200 release of HCP subjects, in [2] S900 release of HCP subjects). Strikingly, the first two Functional Harmonics 1 and 2 reflect the principal gradients of human cortex delineating unimodal and transmodal networks of the brain [3] both in the 7 and 17 RSN implementation [1]. In the 17 RSN implementation, FHs 3 and 4 demonstrate separation of SMN and VIS. Moreover, FH5 places in opposite polarities the SMN and DMN (mainly its temporal parietal part) networks on one side, and DAN and FPN on the other. FH6 separates SMN and the temporal parietal part of the

DMN from the rest of the DMN and FPN. FH 7 further differentiates the VIS and SM. FH 8 splits attention networks (mainly VAN) and DMN. FH 9 demonstrates gradient like separation between DMN and other higher-order cognitive networks (DAN, VAN, LN and FPN). FHs 10 and 11 strongly represent part of the FPN (and LN in the 7 RSNs representation) (**Figure SI 1B**).

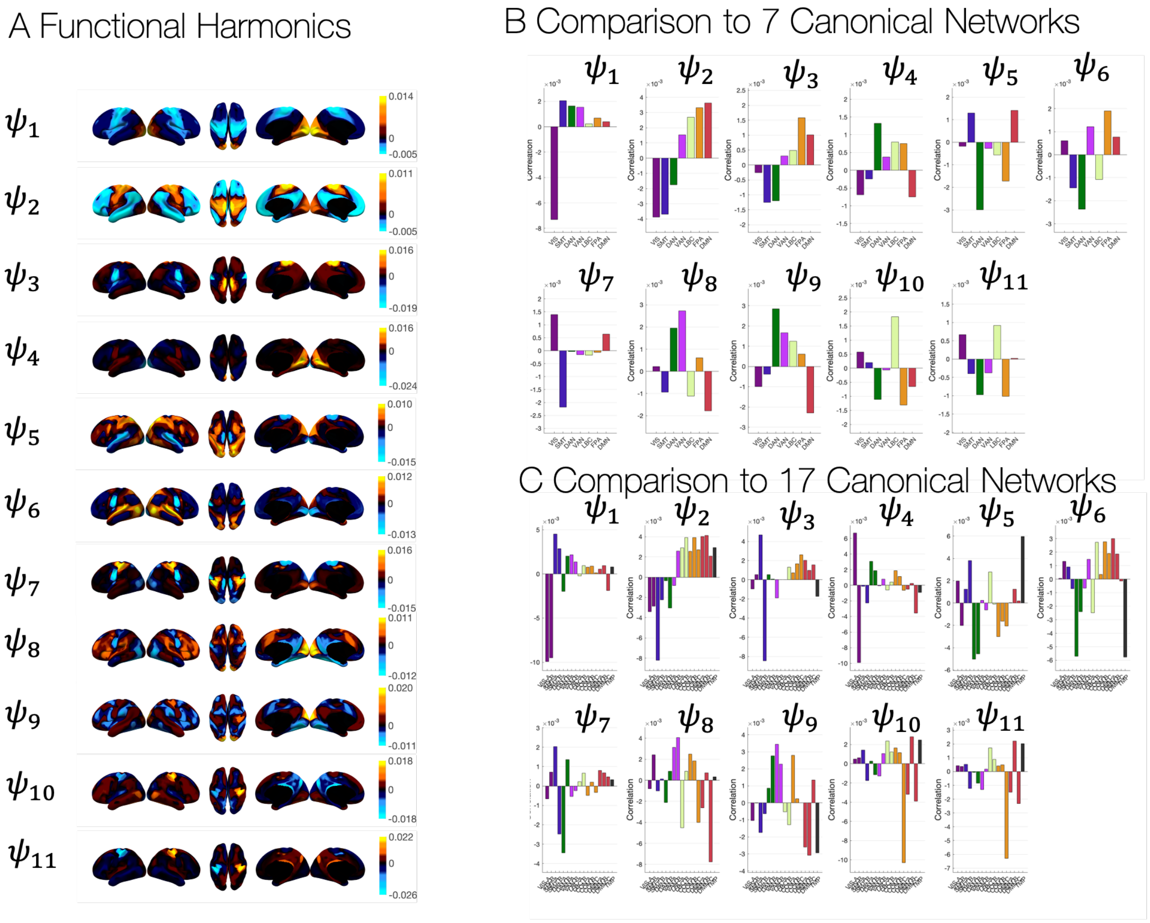


***Figure SI 1: Functional Harmonics: A)*** *The first 11 Functional Harmonics as obtained from the Laplacian eigendecomposition of the dense functional connectome.* ***B)*** *Comparison with 7 canonical resting-state networks.* ***C)*** *Comparison with 17 canonical resting-state networks* [1]. *Different Functional Harmonics indicate separation into various weighted combinations of resting state networks.*

### Functional Harmonic Reconstruction

The reconstructed signal can be further defined as follows

$$\mathcal{F}^R(x, t_i) = \sum_{k=1}^{n} \psi_k(x)\, \tau_k(t_i)$$

where $\mathcal{F}^R(x, t_i)$ is the reconstructed signal obtained from spatial components (Functional Harmonics), $\psi_k(x)$, and temporal components (contributions of Functional Harmonics), $\tau_k(t_i)$, at every timepoint $t_i$ (**Figure 1 E**).

### Signal Reconstruction

By projecting Functional Harmonics onto the timeseries, it is possible to obtain the contribution of each FH evolving in time. Henceforth, the underlying spatio-temporal activity is described in terms of its spatial (FH) and temporal (signal of FH contributions) dimensions and can be reconstructed back by linear summation (**Figure SI 2A**). For a template (DMT pre-treatment) condition, we show that reconstruction with 100 FHs achieves a correlation of 0.5, and 0.6 when the $0^{th}$ global FH is considered, with the original signal. Furthermore, when taking only the first 11 FHs for the reconstruction, the reconstructed signal correlates 0.4 with the original signal and further 0.5 when the 0th global FH is considered (**Figure SI 2B**).

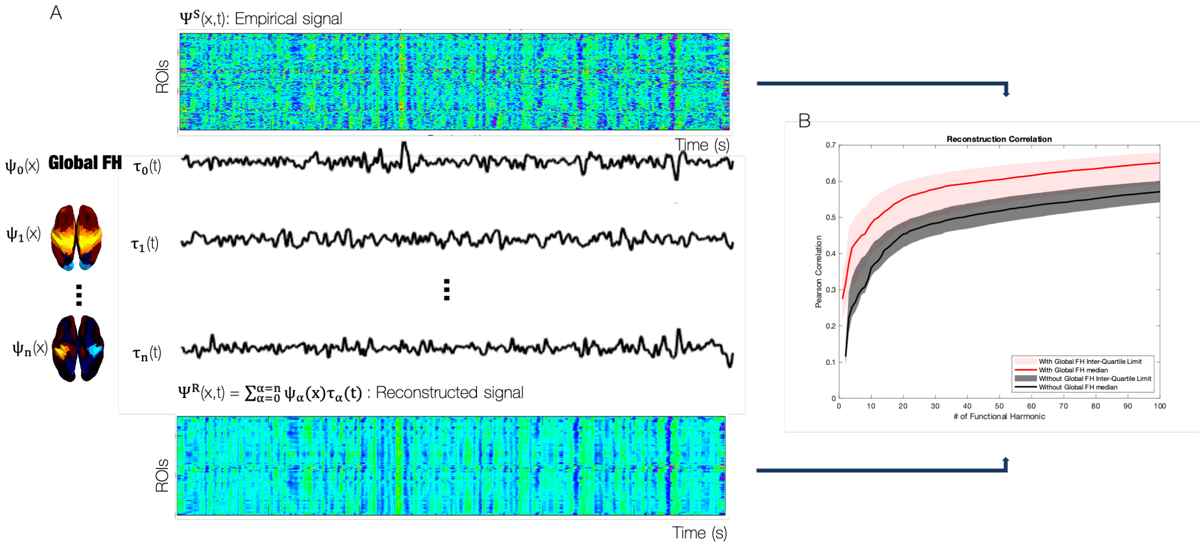


***Figure SI 2: Signal Reconstruction:*** *A) After the Functional Harmonic Decomposition, a signal can be reconstructed back by combining the spatial component (FHs) and temporal component (FH contribution). Furthermore, a good portion of the signal can be reconstructed with only a fraction of the FHs. B) Correlation of reconstruction between the empirical signal of DMT before injection and its reconstructed counterpart with varying number of FHs. The reconstruction with 100 FHs achieves a correlation of 0.5, and 0.6 with the 0$^{th}$ global FH. Furthermore, only when considering the first 11 FHs, they correlate 0.4 and when the 0th global FH is included 0.5.*

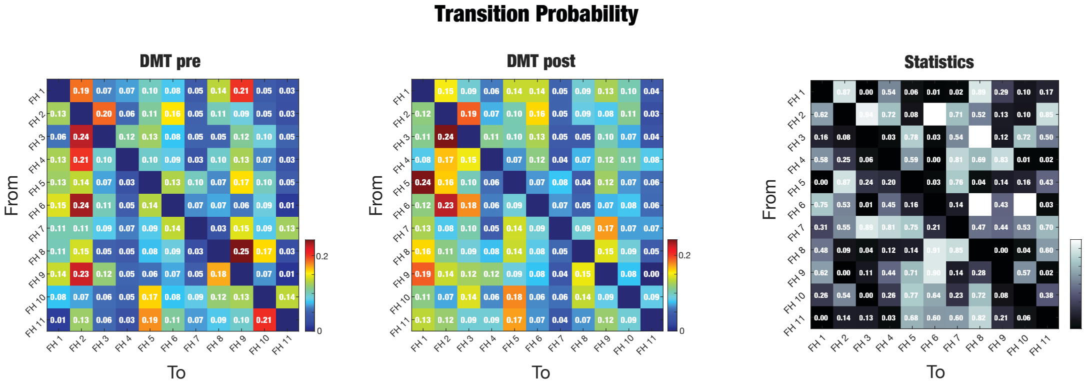


***Figure SI 3: Transition Probability Matrix****. Transition Probability matrices for the two DMT conditions. We also report its statistics (p-value < 0.05 uncorrected paired t-test).*

[1] B. T. T. Yeo *et al.*, “The organization of the human cerebral cortex estimated by

intrinsic functional connectivity.," *J. Neurophysiol.*, vol. 106, no. 3, pp. 1125–65, Sep. 2011.

[2] K. Glomb, M. L. Kringelbach, G. Deco, P. Hagmann, J. Pearson, and S. Atasoy, "Functional harmonics reveal multi-dimensional basis functions underlying cortical organization," *Cell Rep.*, vol. 36, no. 8, 2021.

[3] D. S. Margulies *et al.*, "Situating the default-mode network along a principal gradient of macroscale cortical organization," *Proc. Natl. Acad. Sci. U. S. A.*, vol. 113, no. 44, pp. 12574–12579, Nov. 2016.
